# Extracellular signal-regulated kinase (ERK) pathway control of CD8^+^ T cell differentiation

**DOI:** 10.1101/2020.08.18.255711

**Authors:** Marcos P. Damasio, Julia M. Marchingo, Laura Spinelli, Doreen A. Cantrell, Andrew J.M. Howden

## Abstract

The integration of multiple signalling pathways that co-ordinate T cell metabolism and transcriptional reprogramming is required to drive T cell differentiation and proliferation. One key T cell signalling module is mediated by extracellular signal-regulated kinases (ERKs) which are activated in response to antigen receptor engagement. The activity of ERKs is often used to report antigen receptor occupancy but the full details of how ERKs control T cell activation is not understood. Accordingly, we have used mass spectrometry to explore how ERK signalling pathways control antigen receptor driven proteome restructuring in CD8 ^+^ T cells to gain insights about the biological processes controlled by ERKs in primary lymphocytes. Quantitative analysis of >8000 proteins identified only 900 ERK regulated proteins in activated CD8^+^ T cells. The data identify both positive and negative regulatory roles for ERKs during T cell activation and reveal that ERK signalling primarily controls the repertoire of transcription factors, cytokines and cytokine receptors expressed by activated T cells. The ERKs thus drive the transcriptional reprogramming of activated T cells and the ability of T cells to communicate with external immune cues.

## Introduction

The growth, proliferative expansion and differentiation of T lymphocytes is initiated by signalling pathways regulated by the T cell antigen receptor (TCR) and then balanced by positive and negative feedback signals transduced by cytokines, co-stimulatory receptors and inhibitory/immune checkpoint receptors (1). The initial events in TCR signalling are mediated by cytosolic tyrosine kinases and adaptors that function to couple the TCR to a network of serine-threonine kinases that propagate the signal from the cell membrane to the nucleus and drive the transcriptional and metabolic changes that support effector T cell differentiation (2). One important serine-threonine kinase signalling cascade regulated by the TCR is mediated by the mitogen-activated protein kinases (MAPK) ERK1 and ERK2 (3-5). The activation of ERK1/2 in T cells is controlled by Ras guanine nucleotide–binding proteins. TCR triggering rapidly causes Ras proteins to cycle from a GDP-bound (inactive) to a GTP-bound (active) state that allows Ras proteins to bind to Raf serine-threonine kinases. This drives a pathway whereby active Raf kinases phosphorylate and activate the kinases MEK1/2 which then phosphorylate key threonine and tyrosine residues in ERK1/2 to activate these kinases (5, 6).

The TCR acts as a digital switch for ERK1/2 activation in that the strength of the antigen stimulus determines the frequency of T cells that activate ERKs. This is a very sensitive switch with a low threshold for activation, and even very low levels of antigen receptor occupancy can activate the entire ERK1/2 pool in a T cell (4, 7). Moreover, ERK1/2 signalling is important for TCR function during positive selection in the thymus and in peripheral T cells. For example, activation of ERK2 is important for TCR induced T cell proliferation and controls the survival of TCR activated peripheral CD8^+^ T cells (8). The importance of ERK1/2 activity for T cells and the sensitivity of flow cytometric assays to quantify the phosphorylation and hence the activation of ERK1/2 has promoted the use of this pathway as a sensitive readout of TCR receptor occupancy. However, although ERK1/2 activity is used to assess TCR signalling capacity the full details of how the RAS/MAPK cascade controls T cell function is not clear. Substrates for ERK1 and ERK2, that give some insights as to why this kinase pathway is so important in T cells, include the ternary complex factor subfamily of ETS-domain transcription factors ELK-1, SAP-1 and SAP-2, which control expression of immediate early genes such c-Fos, Egr1 and Egr3 in T cells (9). The Erk-mediated regulation of c-Fos expression contributes to the activator protein-1 (AP-1) transcription complex formed by the transcription factors c-Fos and Jun (10, 11). Egr1 and Egr3 play a critical role in T cell activation, regulating the expression of interleukin 2 (IL-2) and T cell proliferation (12, 13). There are also other ways in which ERK1/2 signalling can control T cell function. For example, the Ras/MAPK pathway can control microtubule remodelling through the phosphorylation and regulation of stathmin (14). Moreover, a number of serine threonine kinases are phosphorylated and activated by the ERKs including the 90-kilodalton ribosomal protein S6 kinases (RSK1 and RSK2) which have been implicated in the control of cell cycle progression and cytokine production in activated T cells (15).

Quantitative analysis of T cell proteomes using high resolution mass spectrometry is increasingly being used to understand T cell biology and provide new insight into how T cells respond to immune challenges and differentiate to effector cells (16-20). The importance of such proteome analysis stems from the impact that changes in the rates of protein production ie rates of protein synthesis versus degradation can have on how a cell executes its transcriptional program. We have shown recently that antigen receptor engagement causes a remodelling of T cell proteomes by increasing and decreasing expression of more than 6,000 and 1,000 proteins respectively (16). Accordingly, the challenge is to understand the regulatory contribution of the different TCR signalling pathways to this proteome remodelling. In this context, we have recently mapped how the metabolic regulators mTORC1 and Myc control the proteome restructuring of immune activated cells (16, 18). The objective of the present report was to explore the ERK1/2 contribution as a control switch for antigen receptor induced proteome remodelling. These data reveal that the ERK1/2 signalling pathway regulates a relatively small fraction of the proteome restructuring program initiated by the TCR. The data also identify both positive and negative regulatory roles for ERK signalling during T cell activation and show moreover that the dominant functions of ERK1/2 in T cells are to control the repertoire of transcription factors, cytokines and cytokine receptors expressed by activated T cells. The ERK1/2 controlled T cell proteins, while relatively small in number, includes many of the key molecules known to be pivotal for T cell differentiation and acquisition of effector function.

## Materials and methods

### Mice

For CD8^+^ TCR activated proteomes P14 (21) transgenic mice were used (female mice, 8-10 weeks old). Naïve CD8^+^ cells used for proteomics were described previously (18). For the analysis of cell markers by flow cytometry 3 x Myc-eGFP (22) female mice aged 12-22 weeks were used. For cell proliferation assays P14 transgenic mice were used (male mice, 9 −10 weeks old). All mice were maintained in the Biological Resource Unit at the University of Dundee using procedures approved by the University Ethical Review Committee and under the authorization of the UK Home Office Animals (Scientific Procedures) Act 1986.

### Cells and flow cytometry

All cells were cultured at 37 °C with 5 % CO^2^ in RPMI 1640 containing glutamine (Invitrogen) and supplemented with 10 % FBS (Gibco), 50 μM β-mercaptoethanol (Sigma) and penicillin/streptomycin (Gibco). For proteomics experiments in vitro TCR stimulation of lymphocytes was performed as follows: lymph nodes from P14 transgenic mice were removed and mashed in RPMI media in a 70 μm cell strainer. Cells were suspended in RPMI media and stimulated for 24 hours with 100 ng/ml antigenic peptide GP33 (glycoprotein amino acids 33–41) and +/-2 μM PD184352. After 24 hours, cells were collected and prepared for cell sorting to generate a pure population of CD8^+^ cells. Harvested cells were treated with 1 μg Fc block (BD Pharmingen) per million cells, to block Fc receptors. Cells were stained with CD8 - PE and DAPI and sorted on an Influx cell sorter (Becton Dickinson). CD8^+^ viable cells were collected and washed with HBSS before being snap frozen in liquid nitrogen. Pure populations of naïve CD8^+^ cells were prepared as described previously (18).

For flow cytometry experiments to examine marker expression, lymph nodes were removed from Myc-eGFP mice and mashed and cell suspension activated with anti-CD3 (1 µg/mL, clone 145-2C11, Biolegend) and anti-CD28 (3 µg/mL, clone 37.51, eBioscience) +/-inhibitor PD184352 (2 µM) at a density of 2 million live cells / mL in a total of 1 mL of RPMI complete medium in a 48 well plate. Cells were placed in culture (37°C, 5% CO^2^) for 24 hrs. After activation cells were harvested by centrifugation and blocked for 5 min with Fc block. Stained with CD4 BV510 (Biolegend RM4-5), CD8 BV421 (Biolegend 53-6.7), CD69 PerCPCy5.5 (Biolegend H1.2F3), CD25 PECy7 (Biolegend PC61), CD62L APC (BD MEL-14), CD44 APCeF780 (eBioscience IM7) all at 1:200 in FACS buffer (DPBS 1%FBS) for 20 min at 4 °C in the dark. Cells were washed with FACS buffer, fixed with IC fix (eBioscience) for 15 min and washed and suspended in FACS buffer before being analysed on a Novocyte flow cytometer.

For cell proliferation assays splenocytes freshly isolated from P14 mice were labelled with 5 μM CFSE (Invitrogen) at 37 °C for 20 min in RPMI complete media (described above). Cells were labelled at a density of 10×10^6^ cells/ml. Excess CFSE was washed off with RPMI and CFSE-labelled cells were seeded at 1⨯10^6^ cell/ml and activated +/-2 μM PD184352 with 100 ng/ml antigenic peptide GP33-41. Unlabelled cells and CFSE-labelled cells incubated in media supplemented with IL-7 were used as controls. Proliferation of CD8^+^ cells was assessed after 48 hours by flow cytometric analysis by monitoring the dilution of CFSE label.

### Proteomics sample preparation and peptide fractionation

Cell pellets were lysed as described previously (16). In brief, cells were lysed in 400 μl lysis buffer (4% sodium dodecyl sulfate, 50 mM tetraethylammonium bromide (pH 8.5) and 10 mM tris(2-carboxyethyl)phosphine-hydrochloride). Lysates were boiled for 5 minutes and sonicated with a BioRuptor (15 cycles: 30 s on and 30 s off) before alkylation with 20 mM iodoacetamide for 1 h at 22 °C in the dark. Cell lysates were subjected to the SP3 procedure for protein clean-up (23) before elution into digest buffer (0.1% sodium dodecyl sulfate, 50 mM tetraethylammonium bromide (pH 8.5) and 1 mM CaCl^2^) and digested with LysC and Trypsin, each at a 1:50 (enzyme:protein) ratio. Peptide clean-up was performed as described in the SP3 protocol (23) and peptides were eluted in 2 % DMSO for fractionation.

Peptides from 24 hour antigen activated cells were fractionated using high pH reverse-phase chromatography as described previously (16). Samples were loaded onto a XbridgeTM BEH130 C18 column with 3.5 μm particles (Waters). Using a Dionex BioRS system, the samples were separated using a 25-min multistep gradient of solvents A (10 mM formate at pH 9 in 2% acetonitrile) and B (10 mM ammonium formate at pH 9 in 80% acetonitrile), at a flow rate of 0.3 ml min−1. Peptides were separated into 16 fractions. The fractions were subsequently dried, and the peptides were dissolved in 5% formic acid and analyzed by liquid chromatography–mass spectrometry. Peptides from naïve cells were fractionated as above but with slight modifications (18).

### Liquid chromatography mass spectrometry analysis (LC-MS/MS)

LC-MSMS was performed as described previously but with slight modifications (24). For each sample, 1 μg of peptides was injected onto a nanoscale C18 reverse-phase chromatography system (UltiMate 3000 RSLC nano, Thermo Scientific) and electrosprayed into an Orbitrap mass spectrometer (LTQ Orbitrap Velos Pro; Thermo Scientific). For chromatography the following buffer conditions were used: HPLC buffer A (0.1% formic acid), HPLC buffer B (80% acetonitrile and 0.08% formic acid) and HPLC buffer C (0.1% formic acid). Peptides were loaded onto an Acclaim PepMap100 nanoViper C18 trap column (100 µm inner diameter, 2 cm; Thermo Scientific) in HPLC buffer C with a constant flow of 10 µl/min. After trap enrichment, peptides were eluted onto an EASY-Spray PepMap RSLC nanoViper, C18, 2 µm, 100Å column (75 µm, 50 cm; Thermo Scientific) using the buffer gradient: 2% B (0 to 6 min), 2% to 35% B (6 to 130 min), 35% to 98% B (130 to 132 min), 98% B (132 to 152 min), 98% to 2% B (152 to 153 min), and equilibrated in 2% B (153 to 170 min) at a flow rate of 0.3 µl/min. The eluting peptide solution was automatically electrosprayed using an EASY-Spray nanoelectrospray ion source at 50°C and a source voltage of 1.9 kV (Thermo Scientific) into the Orbitrap mass spectrometer (LTQ Orbitrap Velos Pro; Thermo Scientific). The mass spectrometer was operated in positive ion mode. Full-scan MS survey spectra (mass/charge ratio, 335 to 1800) in profile mode were acquired in the Orbitrap with a resolution of 60,000. Data were collected using data-dependent acquisition: the 15 most intense peptide ions from the preview scan in the Orbitrap were fragmented by collision-induced dissociation (normalized collision energy, 35%; activation Q, 0.250; activation time, 10 ms) in the LTQ after the accumulation of 5000 ions. Precursor ion charge state screening was enabled, and all unassigned charge states as well as singly charged species were rejected. The lock mass option was enabled for survey scans to improve mass accuracy.

### Proteomics data processing and analysis

Raw mass spec data files were searched using the MaxQuant software package (version 1.6.2.6). Proteins and peptides were identified using a hybrid database from the Uniprot release 2019 07. The hybrid database was generated using all manually annotated mouse SwissProt entries, combined with mouse TrEMBL entries with protein level evidence available and a manually annotated homologue within the human SwissProt database. The following search parameters within MaxQuant were selected: protein N-terminal acetylation and methionine oxidation were set as variable modifications and carbamidomethylation of cysteine residues was selected as a fixed modification; trypsin and LysC were selected as digestion enzymes with up to 2 missed cleavages; the false discovery rate was set at 1 % for protein and peptide and the match between runs function was disabled. Proteins were removed from the data set which were categorised as “reverse”, “contaminant” or “only identified by site”. Estimates of protein copy numbers per cell were calculated using the histone ruler method (25) within the Perseus software package (26).

### Statistics and calculations

To identify significant changes in protein abundance P values were calculated using a two-tailed t-test with unequal variance, on log-normalised protein copy numbers. Proteins were considered to change significantly with a P value <0.05 and a fold change >1.5 or <0.66. To identify processes that may be enriched upon ERK inactivation a more stringent fold change cut-off was established. The standard deviation of the log2(fold change) for all proteins was calculated and a significance cut-off was set as 2 standard deviations from the mean log2(fold change). The mass of individual proteins was estimated using the following formula: CN × MW/NA = protein mass (g cell−1), where CN is the protein copy number, MW is the protein molecular weight (in Da) and NA is Avogadro’s Constant. Heat maps were generated using the Morpheus tool from the Broad Institute (https://software.broadinstitute.org/morpheus). Only proteins with a copy number of at least 500 in all 3 replicates and at least 1 cell population (naïve, TCR activated or TCR activated + inhibitor) were included in the heat map.

## Results

### ERK1/2 selectively remodels the proteome of antigen receptor activated T cells

Quantitative high-resolution mass spectrometry was used to resolve proteomes of CD8^+^ T cells after 24 hours of antigen activation in the presence or absence of the inhibitor PD184352, a highly selective inhibitor of MEK (27), which leads to effective inhibition of T cell antigen receptor induced ERK1/2 activity. These experiments used P14 CD8^+^ T cells which express a TCR specific for lymphocytic choriomeningitis virus glycoprotein peptide gp33-41 (21) and hence allow an assessment of how ERK signalling shapes T cell responses to peptide/MHC complexes. We also compared the proteomes of activated cells to those of naïve CD8^+^ cells, allowing us to evaluate the impact of ERK1/2 activity on TCR induced proteome remodelling. We identified over 8000 proteins and estimated protein copy numbers per cell and protein abundance using the ‘proteomic ruler’ method which uses the histone mass spectrometry signal as an internal standard (25) (Supplementary File 1). Antigen activation of naïve CD8^+^ T cells caused a large increase in total cell protein content (Fig 1A). Interestingly, blocking ERK activity had only a modest effect on the mass of antigen activated cells (Fig 1A), with cells still achieving a large increase in protein content compared with naïve cells. This was supported by flow cytometry data of cell forward and side scatter, which revealed that blocking ERK activity had no significant impact on the estimated size of T cells after 24 hours of TCR activation (Fig 1B). To assess the efficacy of the ERK inhibitory strategy we first looked at the effects of PD184352 on the expression of previously defined targets for ERK signalling pathways in T cells: the adhesion molecule CD69 and the transcription factors EGR1 and EGR2. The data show that the upregulation of CD69 and EGR1 and 2 expression which normally occurs in antigen activated T cells was supressed when ERK activity was blocked (Fig 1C, D and E).

**Figure 1.**
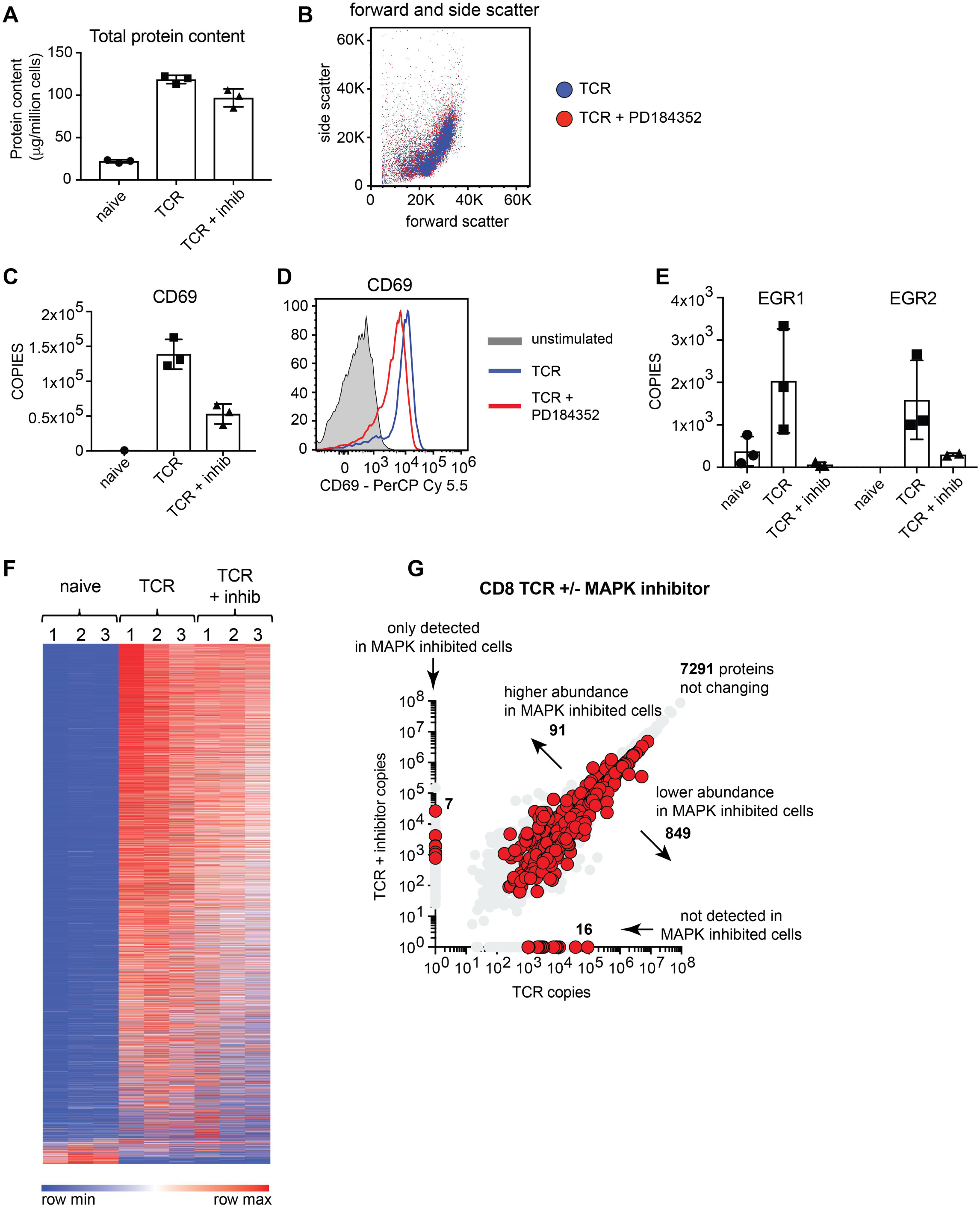
Selective proteome remodelling by ERK. High resolution quantitative mass spectrometry was used to characterise the proteomes of naïve and 24 hour antigen activated CD8^+^ T cells +/-PD184352, a selective inhibitor of ERK activation (TCR and TCR + inhib). (a) Total protein content of T cell populations. (b) Forward/side scatter flow cytometry analysis of control and inhibitor treated cells after 24 hours of antigen activation. (c) Mean protein copy numbers per cell of CD69, estimated using the histone ruler method (25). (d) CD69 expression measured by flow cytometry in naïve and activated cells. (e) Mean protein copy numbers per cell of the transcription factors Early Growth Response 1 and 2 (EGR1 and EGR2). (f) Heat maps of naïve and TCR activated CD8^+^ proteomes +/-PD184352. Abundance is graded from low (blue) to high (red) for each individual protein. (g) Protein copy number comparison for control and inhibitor treated cells. Proteins highlighted in red were significantly different between the two populations (p value <0.05, fold change <0.66 or >1.5, 2-tailed t-test with unequal variance) or were exclusively found in one population at >500 copies per cell and were not detected in the other population. Copy numbers are the mean of 3 biological replicates +/-standard deviation.

We next evaluated the global protein expression profile of CD8^+^ T cells activated with antigen with and without ERK1/2 activity using nearest neighbour analysis and Pearson correlation (Fig 1F). We have previously shown that antigen activation triggers significant proteome remodelling in CD8^+^ T cells with thousands of proteins increasing in abundance while a smaller proportion of proteins drop in abundance in response to activation (16). In the current study we found that antigen activated T cells still underwent a striking proteome remodelling despite ERK activity being blocked (Fig 1F and Supplementary Figure 1). Of the 8000 proteins quantified in activated T cells, over 7000 proteins did not change in abundance when ERK activity was blocked (Fig 1G). However, blocking ERK activity did decrease expression of approximately 800 proteins while almost 100 proteins were found at higher levels when ERK activity was blocked (Fig 1G and Table 1). There were also examples of proteins whose expression was completely absent in one population versus the other (Fig 1G and Table 2). For example, lymphotoxin alpha (LTA), tumour necrosis factor (TNF) and interleukin 2 (IL-2), were detected in 24 hour TCR activated control cells but not detected in ERK inhibited cells, while the cell exhaustion associated protein Thymocyte selection-associated high mobility group box protein TOX was not normally found in antigen receptor activated T cells but was detected when T cells were antigen activated but ERK activation was blocked (Table 2). 10 other transcription factors were also found at higher levels in inhibitor treated cells versus control including Kruppel Like Factor 3 (KLF3) and transcription factor 7 (TCF7 or TCF-1) (Fig 2). In this context, T cell receptor activation results in significant changes in the transcription factor profile of CD8^+^ T cells (16). In this study over 300 proteins annotated as transcription factors were identified and the majority of these did not change significantly in response to ERK inactivation (Fig 2 and Supplementary File 1). For example, T-Box Transcription Factor 21 (T-BET/TBX21) is critical for T cell differentiation and was still significantly upregulated in antigen activated T cells when ERK activity was blocked (Fig 2). However, inhibition of ERKs did cause reduced expression of a number of critical transcription factors that are normally increased in expression in activated CD8^+^ T cells including Interferon Regulatory Factor 8 (IRF8), Eomesodermin (EOMES) and Nuclear Factor Interleukin 3 Regulated (NFIL3). These transcription factors all play key roles controlling T cell differentiation (16, 28-31) and were all significantly reduced in abundance when ERK activity was blocked (Fig 2).

**Table 1.**
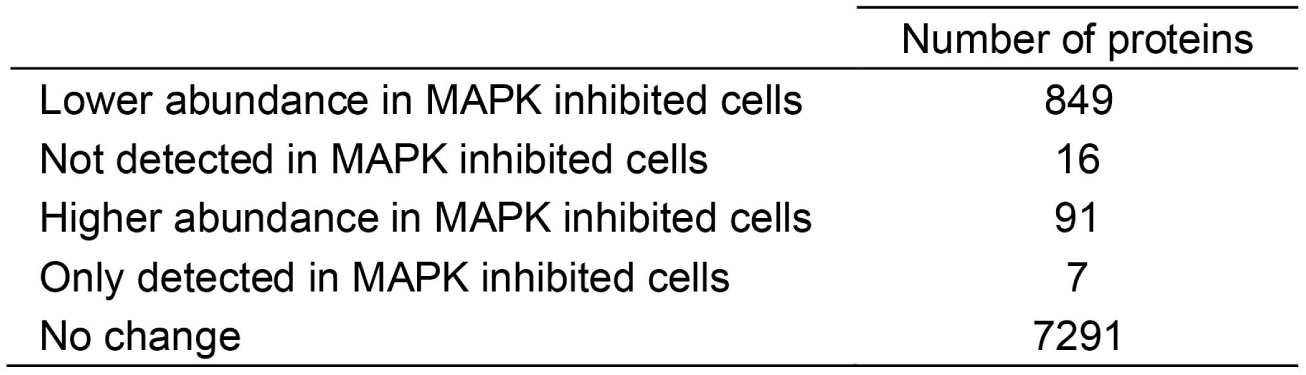
Number of proteins changing in abundance in response to blocking ERK activity. Proteins were considered to change significantly with a p value <0.05 and a fold change <0.66 or >1.5 (2-tailed t-test with unequal variance). Proteins were classified as having a presence/absence expression profile if they were exclusively found at > 500 copies per cell in one population but absent from the other population.

**Table 2.** Proteins showing a presence/absence expression profile in response to blocking ERK activity. Proteins were considered to have a presence/absence expression profile if they were found in all 3 biological replicates of one condition at a copy number > 500 copies per cell and were not detected in any biological replicates of the comparison condition.

| Protein name | Gene name | naïve | TCR | TCR + inhibitor |
| --- | --- | --- | --- | --- |
| RNA-binding protein MEX3D | Mex3d | - | 89,700 | - |
| Lymphotoxin-alpha | Lta | - | 36,100 | - |
| Tumor necrosis factor | Tnf | - | 10,900 | - |
| T-complex protein 1 subunit zeta-2 | Cct6b | 1,100 | 9,100 | - |
| Plasminogen activator inhibitor 1 | Serpine1 | - | 8,400 | - |
| Cytokine-inducible SH2-containing protein | Cish | - | 8,200 | - |
| Runt-related transcription factor 2 | Runx2 | - | 6,900 | - |
| Unconventional myosin-X | Myo10 | - | 3,300 | - |
| Protein BTG3 | Btg3 | - | 3,000 | - |
| Tumor necrosis factor ligand superfamily member 8 | Tnfsf8 | - | 2,600 | - |
| HemK methyltransferase family member 1 | Hemk1 | - | 2,500 | - |
| Mitogen-activated protein kinase kinase kinase 8 | Map3k8 | - | 2,300 | - |
| Interleukin-2 | Il2 | - | 2,200 | - |
| Solute carrier organic anion transporter family member 4A1 | Slco4a1 | - | 2,100 | - |
| Regulator of nonsense transcripts 1 | Upf1 | - | 1,100 | - |
| Psychosine receptor | Gpr65 | - | 1,000 | - |
| Src-like-adaptor 2 | Sla2 | 500 | - | 26,500 |
| Drebrin-like protein | Dbnl | - | - | 4,100 |
| Spindle and kinetochore-associated protein 2 | Ska2 | - | - | 1,900 |
| Peroxisomal membrane protein 4 | Pxmp4 | 600 | - | 1,900 |
| Thymocyte selection-associated high mobility group box protein TOX | Tox | - | - | 1,200 |
| Class II histocompatibility antigen, M alpha chain | H2-DMa | - | - | 1,000 |
| BCL2-like 12 | Bcl2l12 | - | - | 800 |

**Figure 2.**
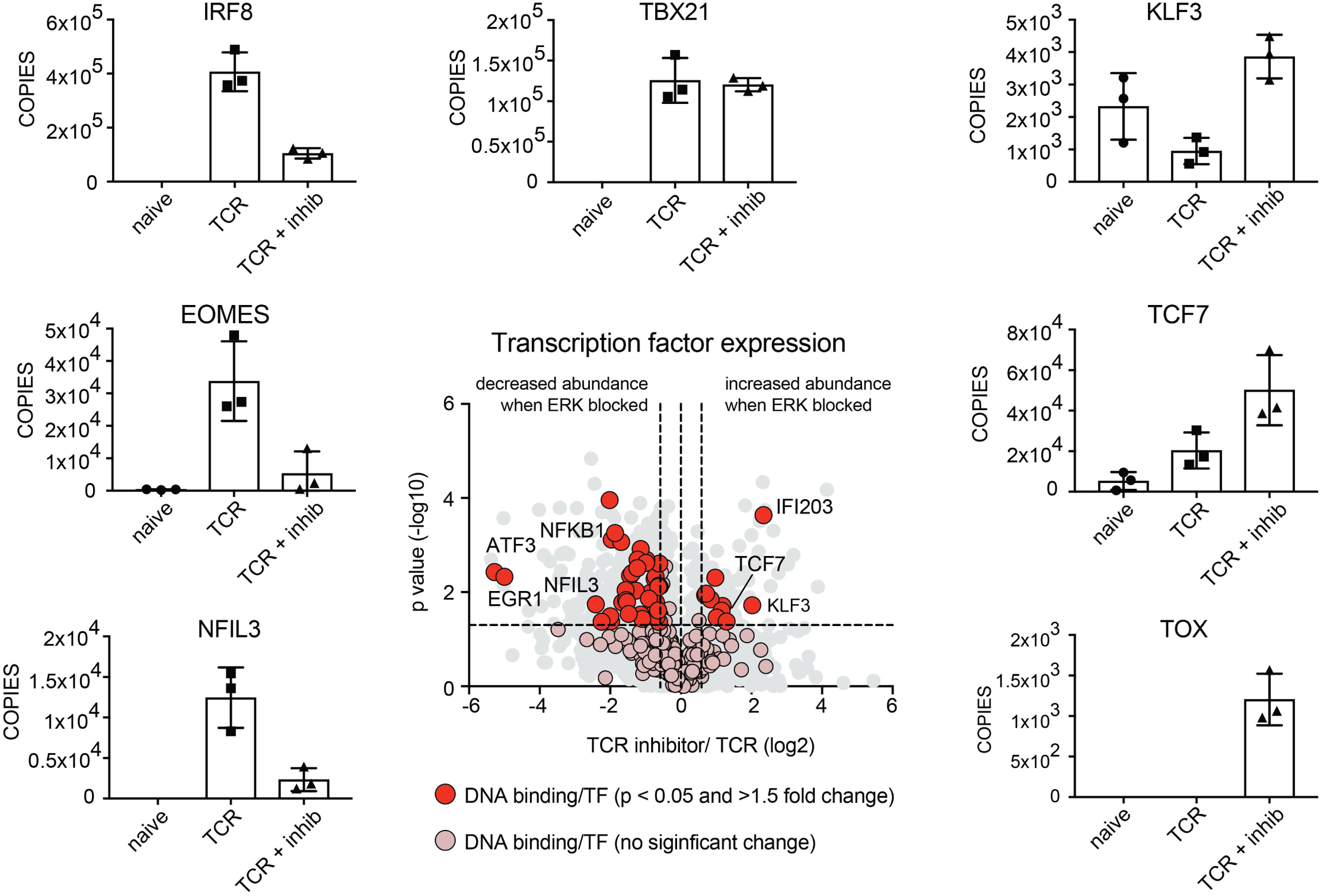
ERK activity is critical for the expression of key transcription factors during CD8^+^ T cell activation. The expression profile of over 300 proteins with the gene ontology term GO:0003700 (DNA binding transcription factor activity) was assessed in response to blocking ERK activity. Proteins highlighted in red were annotated with the above GO term and were significantly different between the two populations (p value <0.05, fold change <0.66 or >1.5, 2-tailed t-test with unequal variance). Proteins highlighted in pink are transcription factors that did not significantly change in response to blocking ERK activity. IRF8, IFN regulatory fa ctor 8; EOMES, eomesodermin; NFIL3, Nuclear Factor Interleukin 3 Regulated; TBX21, T-Box Transcription Factor 21 (T-bet); KLF3, Kruppel Like Factor 3; TCF7, Transcription factor 7; TOX, Thymocyte Selection Associated High Mobility Group Box. Copy numbers are the mean of 3 biological replicates +/-standard deviation.

### ERK1/2 activity controls expression of effector molecules, cytokines and cytokine receptors in activated CD8^+^ T cells

To identify specific biological processes that were impacted by ERK activity we performed a gene ontology enrichment analysis for those proteins that showed the greatest drop in abundance in T cells treated with PD184352 (Table 3). This analysis revealed the top enriched GO term to be “immune response” (GO BP:0006955) which included a number of effector molecules, cytokines and cytokine receptors. An examination of all of the effector molecules identified within the data revealed that granzyme B (GZMB), interferon gamma (IFNG), interleukin 2 (IL-2), lymphotoxin alpha and beta (LTA and LTB), perforin (PRF1) and transforming growth factor beta 1 (TGFB1) were found in lower abundance in inhibitor treated cells compared with control cells (Fig 3A). Other key ‘immune response’ molecules reduced in abundance in ERK inhibited T cells included CD5, CD6, CD200, LAG3 and the integrins ITGA5, ITGAL, ITGAV, ITGB1 and ITGB2 and the Interleukin 4 receptor (IL4R) (Fig 3B). Conversely. the loss of ERK signalling was associated with increased expression of the Interleukin 18 receptor I (Fig 3B and Table 4). The loss of ERK signalling also prevented the downregulation of Interleukin 6 signal transducer (IL6ST) the signalling subunit of the IL-6 receptor. This was expressed at very low levels (300 copies) in control TCR activated cells but was found at more than 4,500 copies in TCR activated cells in which ERK activity had been blocked (Table 4).

**Table 3.** Gene ontology enrichment analysis for those proteins that showed reduced abundance when ERK activity was blocked. Proteins that showed the greatest drop in abundance (p value <0.05 and fold change >2 standard deviations from the mean fold change; two-tailed t-test with unequal variance), were subject to enrichment analysis for biological processes. The top 5 enriched processes are presented along with a list of those proteins identified within the most significant enrichment group “immune response” GO:0006955.

GO term enrichment analysis
| Enriched GO terms for proteins with lower abundance when ERK activity is blocked |  |  |
| --- | --- | --- |
| enriched GO term | fold enrichment | p value |
| GO:0006955~ immune response | 17 | 6.33 x 10<sup>-10</sup> |
| GO:0007165~ signal transduction | 4 | 2.2 x 10<sup>-4</sup> |
| GO:0006959~ humoral immune response | 27 | 3.4 x 10<sup>-4</sup> |
| GO:0010628~ positive regulation of gene expression | 5 | 3.6 x 10<sup>-4</sup> |
| GO:0045080~ positive regulation of chemokine biosynth. | 67 | 7.2 x 10<sup>-4</sup> |

| Immune response GO:0006955 |  |
| --- | --- |
| Gene Name | Protein Name |
| Ifng | Interferon gamma |
| Il2ra | Interleukin 2 receptor, alpha chain |
| Il2 | Interleukin 2 |
| Lta | Lymphotoxin alpha |
| Nfil3 | Nuclear factor, interleukin 3 regulated |
| Serpinb9 | Serine (or cysteine) peptidase inhibitor, clade B, member 9 |
| Tnfsf11 | Tumor necrosis factor (ligand) superfamily, member 11 |
| Tnfsf8 | Tumor necrosis factor (ligand) superfamily, member 8 |
| Tnfrsf18 | Tumor necrosis factor receptor superfamily, member 18 |
| Tnfrsf4 | Tumor necrosis factor receptor superfamily, member 4 |
| Tnf | Tumor necrosis factor |

**Table 4.** The expression profile of cytokine receptors in response to blocking ERK activity. Mean protein copy numbers per cell for cytokine receptors identified within naïve and 24 hour antigen activated CD8^+^ cells +/-PD184352. The fold change (TCR/TCR + inhibitor) is presented along with p value (two-tailed t-test with unequal variance). Proteins highlighted in red showed a significant change in expression when ERK activity was blocked (p value <0.05 and a fold change <0.66 or >1.5).

Cytokine receptors - copies per cell
|  | naïve | TCR | TCR + inhibitor | fold change | P value |
| --- | --- | --- | --- | --- | --- |
| **IL2RA** | - | 103,000 | 18,000 | 0.2 | 0.03 |
| IL2RB | 200 | 10,400 | 12,200 | 1.2 | 0.35 |
| IL2RG | 3,100 | 33,000 | 27,700 | 0.8 | 0.42 |
| **IL4R** | 60 | 3,200 | 500 | 0.2 | 0.00 |
| IL6RA | 700 | - | 200 | - | - |
| **IL6ST** | 900 | 300 | 4,700 | 17.6 | 0.00 |
| IL7R | 2,000 | - | - | - | - |
| IL10RB | - | 2,500 | 1,900 | 0.8 | 0.76 |
| **IL12RB1** | - | 12,700 | 2,100 | 0.2 | 0.01 |
| IL12RB2 | - | 1,800 | 1,800 | 1 | 0.78 |
| IL15RA | - | 5,400 | 3,400 | 0.62 | 0.16 |
| IL17RA | - | 400 | 500 | 1.3 | 0.79 |
| **IL18R1** | 800 | 200 | 3,000 | 12.8 | - |
| IL21R | 400 | 900 | 200 | 0.2 | 0.07 |
| IL27RA | 300 | 1,000 | 1,000 | 1 | - |

**Figure 3.**
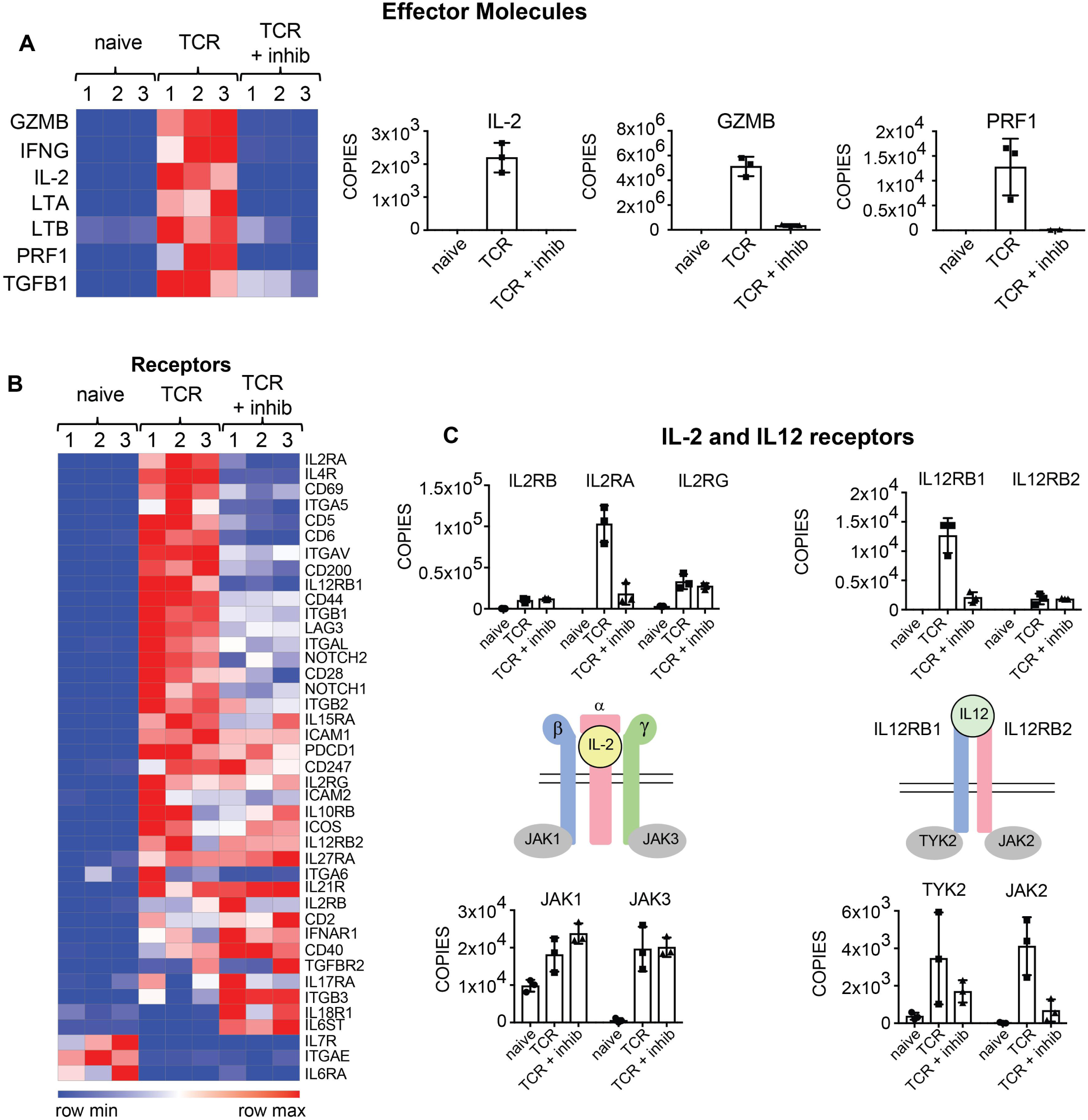
ERK activity controls the expression of effector molecules, cytokine receptors and their downstream signalling components. (a) Expression profile of effector molecules in naïve and antigen activated CD8^+^ T cells +/-PD184352. The heat map shows the relative abundance of individual proteins graded from low (blue) to high (red). GZMB, granzyme B; IFNG, interferon gamma; IL-2, interleukin 2; LTA, lymphotoxin alpha; LTB, lymphotoxin beta; PRF1, perforin 1; TGFB1, Transforming Growth Factor Beta 1. (b) Expression profile of cell surface receptors in naïve and TCR activated CD8^+^ cells +/-PD184352. (c) The abundance of IL-2 and IL-12 receptor subunit components. The IL-2 receptor consists of 3 subunits: IL-2 receptor subunit alpha, beta and gamma (IL2RA, IL2RB and IL2RG), while the IL12 receptor consists of 2 subunits: IL12 receptor subunit beta 1 and 2 (IL12RB1 and IL12RB2). JAK, Janus kinase. TYK2, Tyrosine Kinase 2. Copy numbers are the mean of 3 biological replicates +/-standard deviation.

Two key cytokines for CD8^+^ T cell differentiation are IL-2 and IL-12 which signal through multi subunit receptors that couple to the Janus family tyrosine kinases. The high affinity IL-2 receptor comprises three subunits IL2Rα (CD25), IL2Rβ and IL2Rγ, whereas the IL-12 receptor is a dimer of IL12RB1 and IL12RB2. The IL-2 receptor signals via JAK1 which binds to IL2Rβ and JAK3 which binds to IL2Rγ. The IL-12 receptor signals through TYK2 and JAK2 which bind respectively to IL12RB1 and IL12RB2. In this context, it was striking that blocking ERK signalling severely blunted the normal upregulation of the expression of IL2Rα: this cytokine receptor subunit was not detected in naive T cells, expressed at 100,000 copies in TCR activated cells but only 18,000 copies in ERK inhibitor treated cells (Table 4 and Fig 3C). ERK also controlled the expression of the IL12RB1 subunit which was not detected in naive T cells, expressed at almost 13,000 copies in TCR activated cells but only 2,000 copies in ERK inhibitor treated cells (Table 4 and Fig 3C). However, to understand fully how ERK signalling impacts the cytokine responsiveness of T cells it is important to consider how ERK signalling effects all receptor subunits and their associated tyrosine kinases. In this this context one new insight from the proteomic data is that antigen receptor engagement controls expression of the key tyrosine kinases needed for signal transduction by the IL-2 and IL-12 receptors. Naive T cells were found to constitutively express relatively high levels of JAK1 (approximately 10,000 copies per cell) but only very low levels of JAK2, JAK3 and TYK2 (approximately 50, 500 and 400 copies respectively). The expression of JAK1 increased approximately 2-fold in immune activated T cells whereas increases in expression of JAK2, JAK3 and TYK2 were much more striking, increasing between 10 and 100-fold (Fig 3C). Interestingly, ERK signalling did not equally mediate antigen receptor control of the expression of the different cytokine receptor subunits and JAKs. For example, ERK activation was required for antigen receptor upregulation of IL12RB1, TYK2 and JAK2 but not needed for the expression of IL12RB2 (Fig 3C). There was also no ERK requirement for antigen receptor induced increases in the expression of IL2Rβ, IL2Rγ, JAK1 or JAK3. These results provide the insight that integration of multiple signalling pathways is needed for something as simple as the upregulation of individual cytokine receptor complexes and associated signalling molecules.

### ERKs are not the dominant regulators of T cell metabolic and biosynthetic programs

Regulated changes in T cell metabolism are essential for T cell function (32, 33) so it was important to evaluate whether critical cellular metabolic compartments and processes are regulated by ERK signalling pathway. Ribosomes, glycolytic enzymes and mitochondria make up a larger proportion of the mass of an activated T cell versus a naïve T cell: 5%, 2% and 10% in a naïve cell versus 8%, 5% and 17% in a TCR activated cell (Fig 4A). The increase in cellular protein mass devoted to these processes highlights the shift in the metabolic demand of T cells during activation. ERK activity was not required for this re-modelling of the T cell proteome and T cells with ERK activity blocked still dramatically increased their ribosome, glycolysis and mitochondrial protein mass (Fig 4A). We also looked at how the inhibition of ERKs impacted the regulated expression of amino acid and glucose transporters that deliver the key nutrients that fuel T cell metabolic processes. In this context, we have shown recently that T cells that fail to upregulate the System L amino acid transporter SLC7A5 remain small and have lower expression of >3000 proteins (18). The data in Fig 4B show that in the absence of ERK activation, antigen receptor activated T cells still increase expression of essential amino acid, glucose and lactate transporters. In the absence of ERK signalling the increases in SLC7A5, SLC1A5 (neutral amino acid transporter) and SLC2A1 and SLC2A3 (glucose transporters), were blunted and approximately 2-fold lower than the increases seen in control activated cells (Fig 4C). However, given that we saw no major impact of ERK inhibition on cell mass (Fig 1A) we infer that this modest decrease in expression of the nutrient transporters was not limiting for T cell growth.

**Figure 4.**
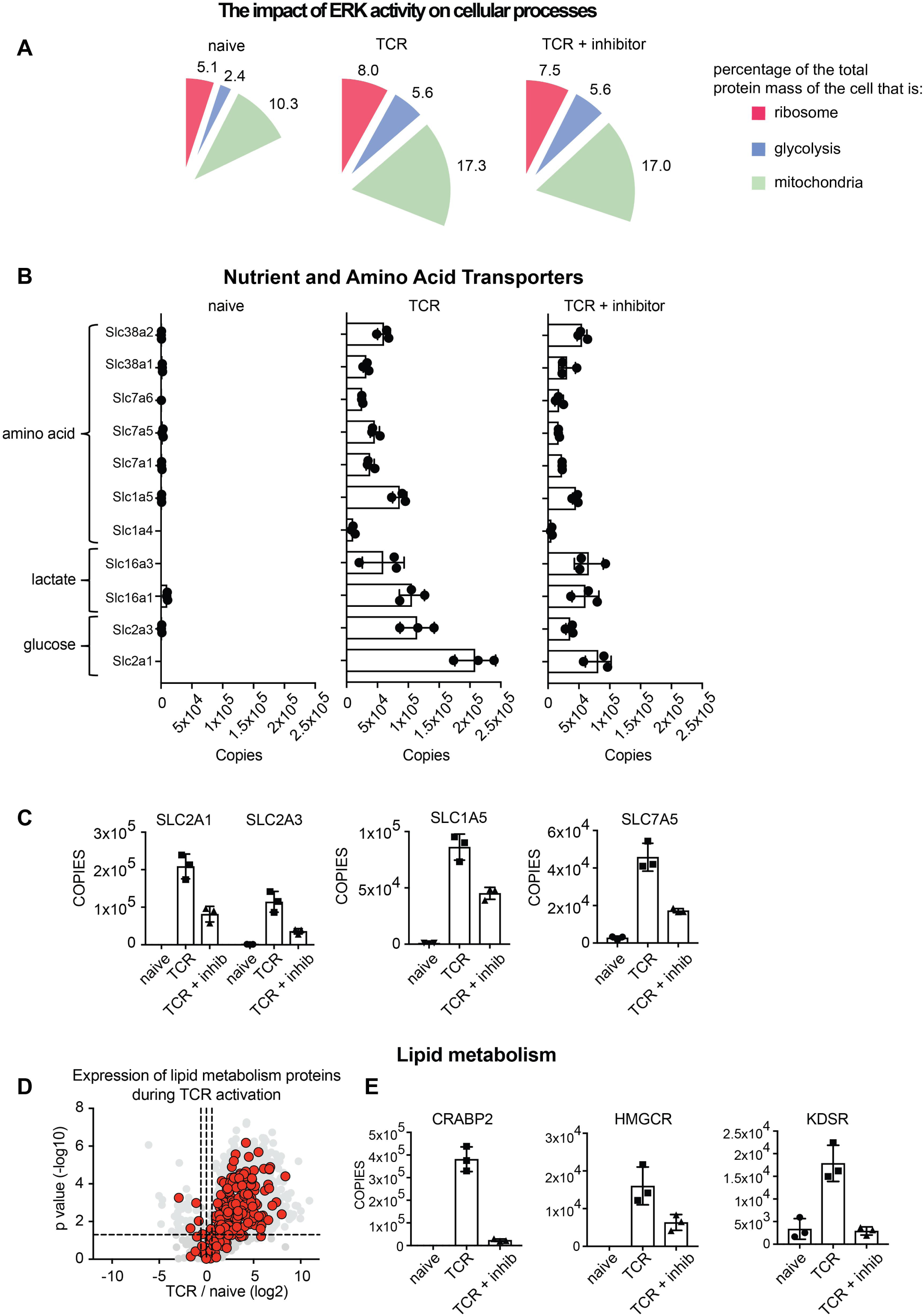
The impact of ERK activity on metabolic processes. (a) Percentage of the total cellular protein mass that represents ribosomal, glycolytic or mitochondrial proteins. (b and c) Expression profile of the major transporters for amino acids, lactate and glucose. (d) The impact of antigen activation on the expression of proteins linked to lipid metabolic processes (GO:0006629). Volcano plot shows the ratio for 24 hour antigen activated cells versus naïve cells. The horizontal dashed line indicates a p value of 0.05 while the outer vertical dashed lines indicate a fold change of 1.5. (e) The mean copy number per cell for a selection of lipid metabolic proteins which were significantly impacted when ERK activity was blocked: CRABP2, Cellular Retinoic Acid Binding Protein 2; HMGCR, 3-Hydroxy-3-Methylglutaryl-CoA Reductase; KDSR, 3-Ketodihydrosphingosine Reductase. Copy numbers are the mean of 3 biological replicates +/-standard deviation.

We also explored the impact of ERK activity on lipid metabolism by examining the expression profiles of proteins that mediate lipid metabolic processes in naive and TCR activated T cells and in T cells activated but with ERK activity blocked (Fig 4D and 4E). We identified over 400 proteins which were annotated as being linked to lipid metabolic processes (GO:0006629). Lipid metabolism was found to be highly TCR regulated with most proteins increasing significantly when naïve CD8^+^ T cells were antigen activated (Fig 4D). Indeed, some lipid metabolism proteins increased >100 fold upon TCR activation including fatty acyl-CoA reductase 1 (FAR1) which increased from 500 copies to 150,000 copies and isopentenyl-diphosphate delta-isomerase 1 (IDI1) which increased from 4,000 copies to over 1.3 million copies when naïve CD8^+^ cells were TCR triggered (Supplementary File 1). Blocking ERK activation impacted the expression of a number of lipid metabolism proteins including cellular retinoic acid binding protein 2 (CRABP2), hydroxymethylglutaryl-CoA reductase (HMGCR) and 3-ketodihydrosphingosine reductase (KDSR) (Fig 4E). The drop-in abundance of CRABP2 when ERK activation was blocked was especially striking, falling from almost 400,000 copies in control TCR activated cells to just over 20,000 copies in ERK inhibited cells. CRABP2 is a retinoic acid (RA) transporter which in turn controls the expression of RA target genes. CRABP2 and RA are key regulators of the differentiation and activity of immune cells (34). Nevertheless, the key conclusion overall from these analyses is that ERKs are not the dominant regulators of T cell metabolic and biosynthetic programs.

### ERK control of CD8^+^ T cell proliferation and survival

Previous studies have shown that the ERK signalling pathway is critical for T cell proliferative expansion (8). The present data reveal that immune activation induced an increase in quantities of three pro-survival BCL2 proteins, MCL1, BCL2 and BCL2L1 and that these were expressed in relatively equal levels in TCR activated T cells (Fig 5). In the absence of ERK signalling the expression of all three of these pro-survival proteins decreased but most notably for BCL2L1 (Fig 5). The data also show that TCR triggering induces increased expression of the pro-apoptotic protein BID and in the absence of ERK signalling the expression of this protein increased. Another pro-apoptotic protein BCL2L11 (BIM) also increased significantly when ERK activation was blocked (Fig 5). D’Souza et al showed previously that ERK signalling controlled expression of Bcl2, Bclx, and Bim mRNA (8). The present proteomic data thus provide strong evidence to support the hypothesis that a key role for ERK signalling is to control T cell survival by controlling the expression of the key proteins that mediate T cell survival.

**Figure 5.**
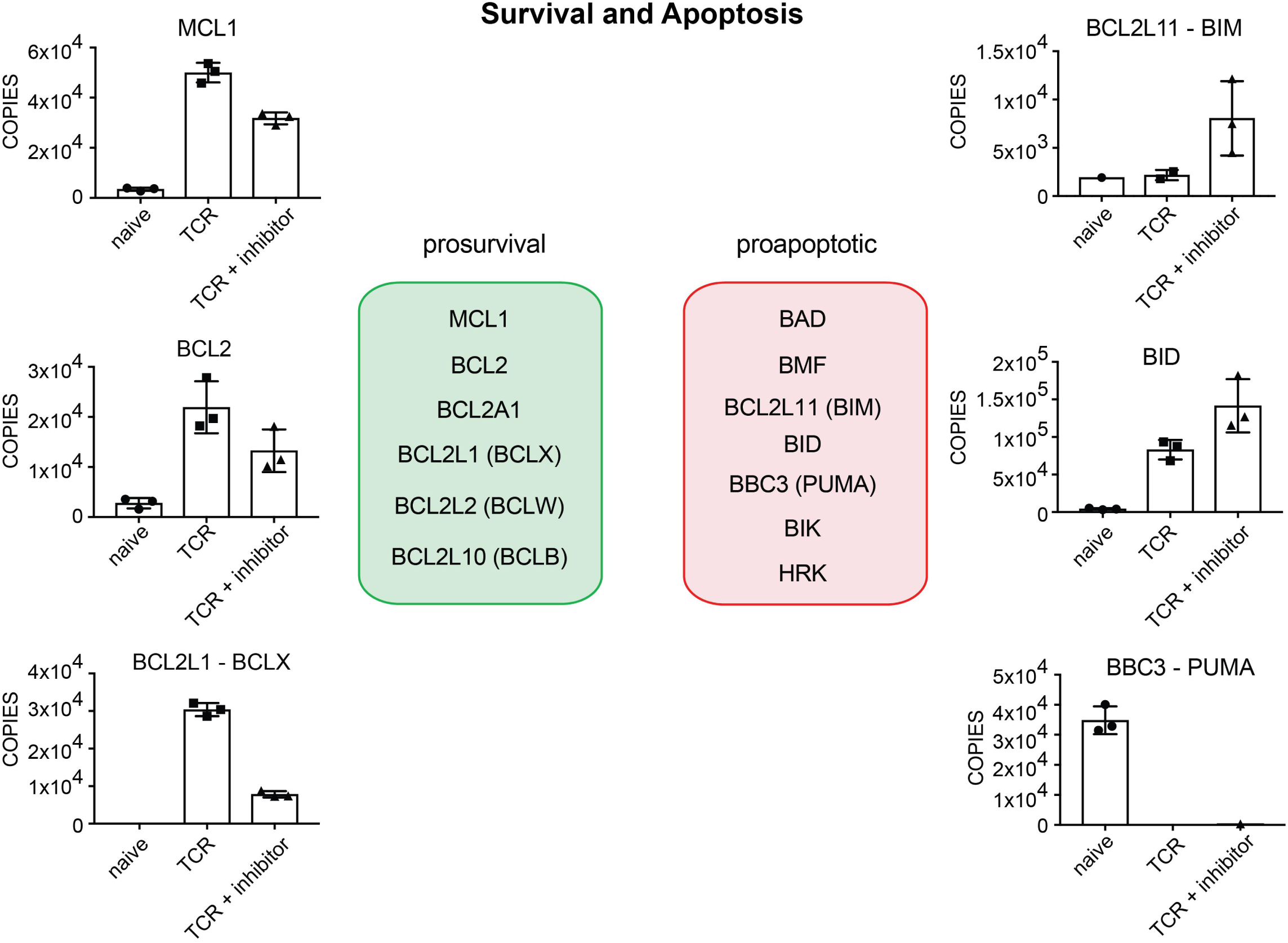
Blocking ERK activity directs cells towards an apoptotic profile. The expression profile of key prosurvival and proapoptotic proteins was assessed in response to blocking ERK activity. MCL1, MCL1 apoptosis regulator BCL2 family member; BCL2, BCL2 apoptosis regulator; BCL2L1 (BCLX), BCL2 Like 1; BCL2L11 (BIM), BCL2 like 11; BID, BH3 Interacting Domain Death Agonist; BBC3 (PUMA), BCL2 Binding Component 3. Copy numbers are the mean of 3 biological replicates +/-standard deviation.

We also interrogated the proteomic data to assess the role of ERK signalling in controlling expression of key cell cycle regulatory proteins. Blocking ERK activity delays CD8^+^ T cell proliferation (Fig 6A). ERK control of cyclin D1 expression is proposed to be important for proliferation in fibroblasts(35). However, T cells do not express cyclin D1, but rather cyclins D2 and 3 dominate and are key for cell cycle progression in T cells (16). T cell activation induces increased expression of cyclin D2, cyclin D3 and their associated kinases CDK4 and CDK6 (Fig 6B) and induces expression of the critical components of the DNA replication fork complex (Fig 6C). GO term enrichment analysis did not identify cell cycle or DNA replication as being enriched in response to ERK inhibitor treatment (Table 3). Indeed, most components of the DNA replication fork complex were not significantly impacted when ERK activity was blocked with the exception of DNA Replication Helicase/Nuclease 2 (DNA2) and DNA polymerase delta subunits 1 and 2 (POLD1 and POLD2) which were found at approximately 50 % lower levels compared to control TCR activated cells (Fig 6C). DNA2, POLD1 and POLD2 are critical enzymes in DNA replication and repair. There was decreased expression of cyclin D2 (CCND2) although cyclin D3 (CCND3) levels were unaffected and cells still had an excess of D type cyclins compared to levels of the cell cycle inhibitor P27 (Supplementary File 1). There was however decreased expression of cyclin E2 (CCNE2) another G1 cyclin. In addition, blocking ERK activity was found to trigger an increase in the expression of Checkpoint Kinase 2 (CHEK2), from just a few hundred copies to over 1,000 copies (Fig 6B). CHEK2 is a DNA damage response protein and cell cycle checkpoint regulator which when activated can arrest the progression of cell cycle, preventing entry to G1 (36). These changes afford an explanation for the ERK signalling requirement for optimal T cell proliferative expansion. However, again, a salient point is that both ERK dependent and independent pathways control expression of the critical machinery needed for DNA replication and T cell cycle progression.

**Figure 6.**
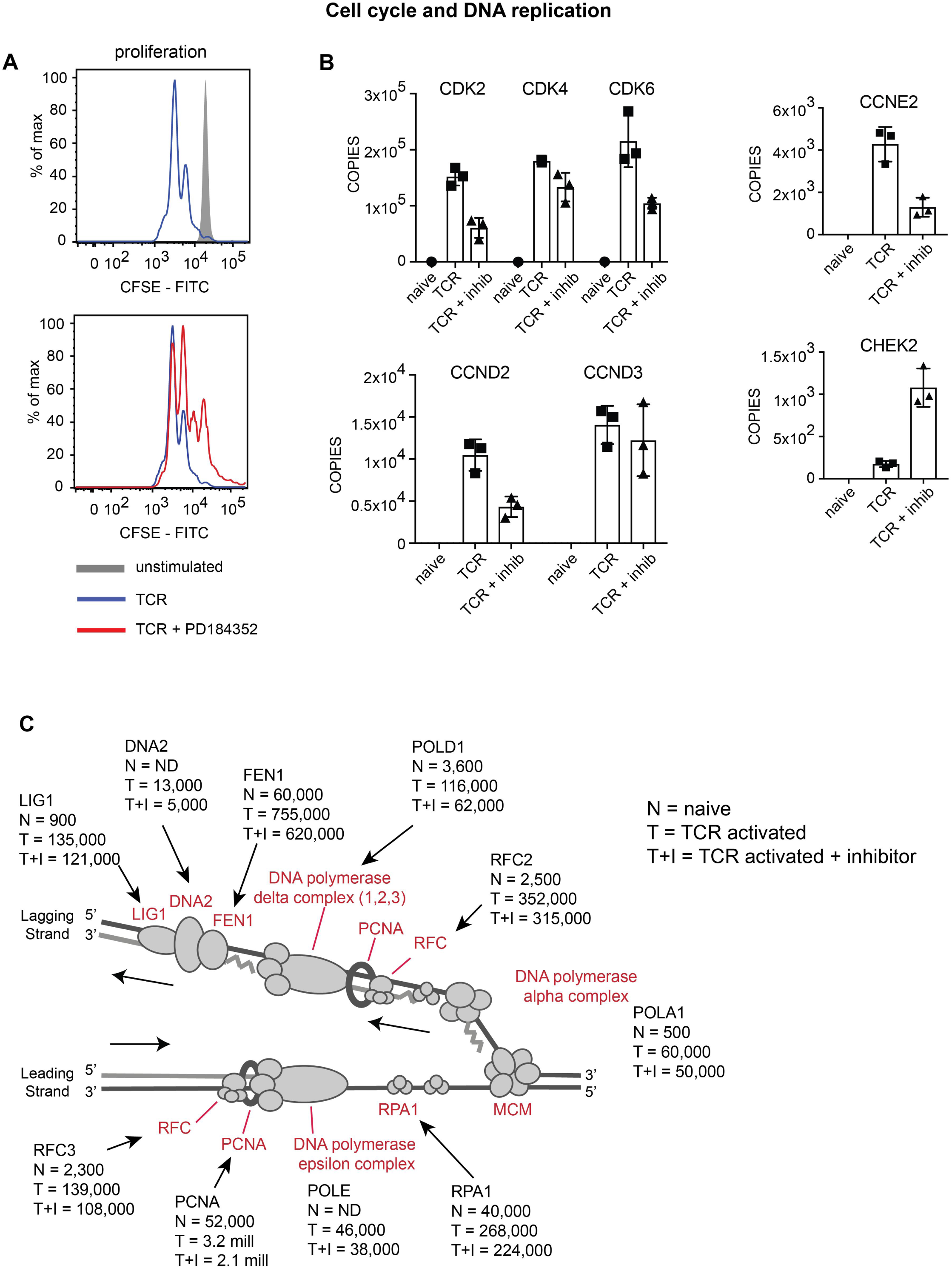
ERK activity has a selective impact on cell cycle and DNA replication machinery. (a) Proliferation of CD8^+^ T cells was assessed after 48 hours of antigen activation by flow cytometric analysis of CFSE label fluorescence. (b) The impact of ERK activity on key cell cycle proteins. CDK2, 4 and 6, cyclin dependent kinase 2, 4 and 6; CCND2 and 3, cyclin D2 and cyclin D3; CCNE2, cyclin E2; CHEK2, checkpoint kinase 2. (c) Expression of components of the DNA replication fork complex in naïve (N), TCR activated (T) and TCR activated + inhibitor (T+I) cells.

## Discussion

The current study has mapped how ERK1/2 regulate antigen driven proteome remodelling of CD8^+^ T cells to understand how these evolutionarily conserved kinases control T cell differentiation. The data show that a large proportion of the proteome restructuring that is driven by triggering of the T cell antigen receptor is not dependent on ERK activation. However, the ERKs are necessary for the production of critical cytokines and cytokine receptors by activated T cells and also control the expression of key transcription factors that drive T cell differentiation. For example, IRF8 integrates antigen receptor and cytokine signals to drive CD8^+^ T cell differentiation (28) and its expression is ERK dependent. Similarly, the expression of NFIL3, which controls the expression of perforin in CD8^+^ T cells (29), requires ERK activation. It was also striking that inhibition of ERKs caused activated T cells to increase expression of the transcription factors TCF7/TCF1 and TOX which are associated with commitment of CD8^+^ T cells to a transcriptional program associated with an exhausted cell phenotype (37-39). The combined effects of ERK inhibition on the expression of positive transcriptional regulators of CD8^+^ T cell differentiation and transcription regulators that drive T cell quiescence/exhaustion programs affords some explanations for why inhibition of ERKs impacts T cell differentiation.

In this context it was intriguing to observe selectivity in the requirements of ERK signalling for the expression of critical components of cytokine receptor complexes. ERK signalling pathways thus do not always have equivalent importance for the expression of different cytokine receptor subunits. This selectivity highlights the importance of understanding whether downregulation of a single subunit of a receptor complex is predictive of loss of receptor function. To answer this question, one needs to have quantitative data that permits an understanding of receptor subunit stoichiometry and then consider what is rate limiting for the production of a high affinity cytokine receptor complex. For example, naive T cells express IL2Rβ and IL2Rγ subunits and JAK1 but have very low levels of JAK3 and no IL2Rα chain. Hence naive CD8^+^ T cells do not have the molecules needed to form a high affinity IL-2 receptor. In response to antigen receptor engagement T cells induce expression of high levels of IL2Rα and also increase abundance of IL2Rβ, IL2Rγ and the essential tyrosine kinases JAK1 and JAK3. It is however valuable to see from the numbers that the limiting IL-2 receptor subunit for formation of a high affinity IL-2 receptors in antigen activated CD8^+^ T cells will be IL2Rβ which is expressed at approximately 10,000 copies per cell. In comparison the activated T cells express more than 100,000 copies of IL2Rα, 30,000 copies of IL2Rγ and approximately 20,000 copies of JAK1 and JAK3. In the absence of ERK signalling, IL2Rα chain abundance reduced more than 5-fold to 18,000 copies per cell but importantly there was no ERK requirement for antigen receptor induced increases in the abundance of IL2Rβ, IL2Rγ, JAK1 or JAK3. Hence, in the ERK inhibited T cells IL2Rα numbers were still in excess of the limiting IL2Rβ subunit allowing the cells to form a high affinity IL-2 receptor complex. In this respect, it is quite common to use flow cytometry analysis of expression of IL2Rα/CD25 to monitor IL-2 responsiveness of T cells. Flow cytometry uses high levels of signal amplification by fluorophore coupled antibodies to create sensitive assays of relative levels of expression of proteins but does not give information about receptor numbers. A 5-fold drop in IL2Rα expression by flow cytometry would look impressive but the current discussion highlights that it is necessary to understand how many copies of IL2Rα are present to understand the consequences of any reductions. Indeed, another insight from quantitative T cell proteomic data is that immune activation also controls the ability of cells to respond to cytokines by controlling expression of the JAK family tyrosine kinases which are integral for cytokine receptor function. Thus, of the 4 members of the JAK family, only JAK1 is expressed at relatively high levels in naive T cells, whereas the other family members, JAK2, JAK3 and TYK2 are very low abundance. T cell activation causes increased expression of all JAK family members and, relevant to the role of ERKs in T cells, the induced expression of JAK2 and TYK2 was dependent on ERK activation. JAK2 activity is necessary for T cell differentiation (40) and this ERK requirement for JAK2 expression would also contribute to ERK control of T cell differentiation.

Finally, a key insight from the present study was that much of the restructuring of the T cell proteome that accompanies T cell activation was ERK1/2 independent. Only 900 of the >8000 proteins quantified in activated T cells were regulated by ERK signalling. In comparison we have shown recently that signalling pathways controlled by Myc and mammalian target of rapamycin complex 1 (mTORC1) controlled expression of more than 4,000 and 2,300 proteins respectively in TCR triggered CD8^+^ T cells (16, 18). There were some points of overlap for Myc, mTORC1 and ERK signalling. For example, they all control expression of cell cycle proteins. However, there were also unique targets for each pathway. A simple example is that ERK but not mTORC1 or Myc signalling controls expression of CD69. However, maximal expression of glucose transporters SLC2A1 and SLC2A3 is sensitive to both the ERK and the mTORC1 pathway but not to Myc (16, 18). In this respect, a key role for Myc is to control expression of amino acid transporters in immune activated T cells and hence in the absence of Myc T cells do not increase cell mass in response to immune activation (18). In contrast, inhibition of ERK signalling has only a very modest effect on the cell mass of an activated T cell. This reflects that ERK mediated signalling is not required to control expression of the most abundant protein groups namely ribosomal, glycolytic and mitochondrial proteins. A comparison of how ERKs, Myc and mTORC1 control T cell proteomes thus provides new insight into how signal transduction pathways guided by the T cell antigen receptor co-ordinate to control T cell differentiation. The major regulators of the key metabolic reprogramming needed for T cell activation identified to date are Myc and mTORC1 (41, 42). However, these metabolic programs allow the T cell to execute its transcriptional program. The key insight from the present study is that ERKs control the transcription factor repertoire of activated T cells and by controlling the production of cytokines and cytokine receptors will dictate the ability of T cells to communicate externally with other cells of the immune system.

## Data Availability

All proteomics data is provided in Supplementary File 1. Raw mass spec data files and MaxQuant analysis files are available from the ProteomeXchange data repository (http://proteomecentral.proteomexchange.org/cgi/GetDataset) and can be accessed with the identifier PXD017147 (for reviewer access - Username, Password: sD4BBgvM. Flow cytometry data are available from the corresponding author upon request.

## Acknowledgments

The authors would like to thank members of the Cantrell lab for comments on the manuscript and M. Lee and R. Clarke from the flow cytometry facility for cell sorting and advice on flow cytometry experiments and J. Hukelmann for performing peptide fractionation. We would also like to thank the biological sciences research unit at the University of Dundee for support with animal work.

## Author Contributions

MD, JMM, LS and AJMH performed experiments. MD and AJMH analysed the proteomics data. MD, AJMH and DAC interpreted the data and AJMH and DAC wrote the manuscript.

---

**Supplementary Figure 1.**
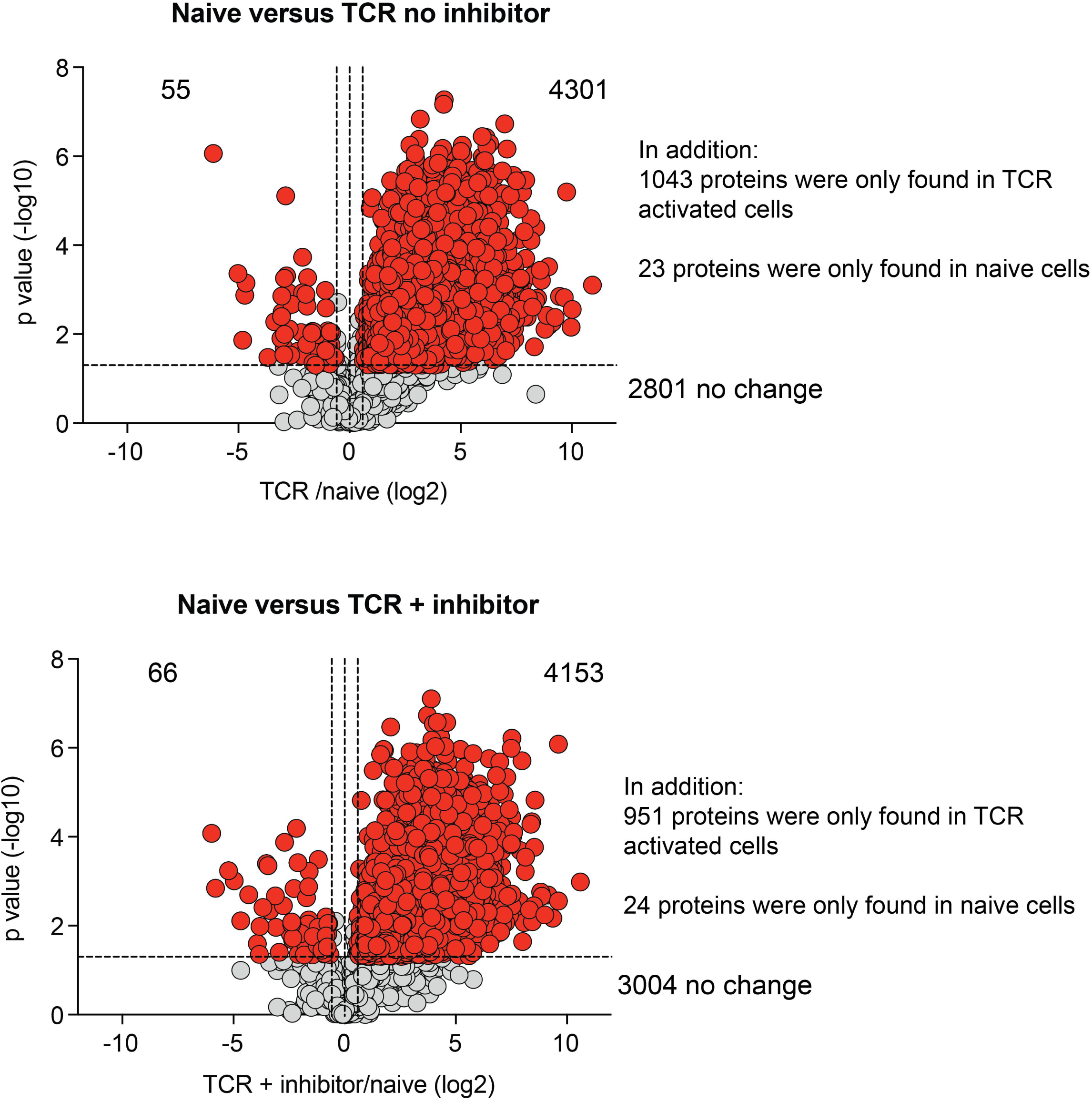

## References

1. Cantrell D. Signaling in lymphocyte activation. Cold Spring Harbor perspectives in biology. 2015;7(6).

2. Navarro MN, Cantrell DA. Serine-threonine kinases in TCR signaling. Nat Immunol. 2014;15(9):808–14.

3. Stefanová I, Hemmer B, Vergelli M, Martin R, Biddison WE, Germain RN. TCR ligand discrimination is enforced by competing ERK positive and SHP-1 negative feedback pathways. Nat Immunol. 2003;4(3):248–54.

4. Altan-Bonnet G, Germain RN. Modeling T cell antigen discrimination based on feedback control of digital ERK responses. PLoS biology. 2005;3(11):e356.

5. Das J, Ho M, Zikherman J, Govern C, Yang M, Weiss A, et al. Digital signaling and hysteresis characterize ras activation in lymphoid cells. Cell. 2009;136(2):337–51.

6. Izquierdo M, Leevers SJ, Marshall CJ, Cantrell D. p21ras couples the T cell antigen receptor to extracellular signal-regulated kinase 2 in T lymphocytes. J Exp Med. 1993;178(4):1199–208.

7. Navarro MN, Feijoo-Carnero C, Arandilla AG, Trost M, Cantrell DA. Protein kinase D2 is a digital amplifier of T cell receptor-stimulated diacylglycerol signaling in naïve CD8^+^ T cells. Sci Signal. 2014;7(348):ra99.

8. D’Souza WN, Chang CF, Fischer AM, Li M, Hedrick SM. The Erk2 MAPK regulates CD8 T cell proliferation and survival. J Immunol. 2008;181(11):7617–29.

9. Rubinfeld H, Seger R. The ERK cascade: a prototype of MAPK signaling. Molecular biotechnology. 2005;31(2):151–74.

10. Karin M, Liu Z, Zandi E. AP-1 function and regulation. Current opinion in cell biology. 1997;9(2):240–6.

11. Treisman R. Regulation of transcription by MAP kinase cascades. Current opinion in cell biology. 1996;8(2):205–15.

12. Collins S, Wolfraim LA, Drake CG, Horton MR, Powell JD. Cutting Edge: TCR-induced NAB2 enhances T cell function by coactivating IL-2 transcription. J Immunol. 2006;177(12):8301–5.

13. Li S, Miao T, Sebastian M, Bhullar P, Ghaffari E, Liu M, et al. The transcription factors Egr2 and Egr3 are essential for the control of inflammation and antigen-induced proliferation of B and T cells. Immunity. 2012;37(4):685–96.

14. Marklund U, Brattsand G, Shingler V, Gullberg M. Serine 25 of oncoprotein 18 is a major cytosolic target for the mitogen-activated protein kinase. J Biol Chem. 1993;268(20):15039–47.

15. Lin JX, Spolski R, Leonard WJ. Critical role for Rsk2 in T-lymphocyte activation. Blood. 2008;111(2):525–33.

16. Howden AJM, Hukelmann JL, Brenes A, Spinelli L, Sinclair LV, Lamond AI, et al. Quantitative analysis of T cell proteomes and environmental sensors during T cell differentiation. Nat Immunol. 2019;20(11):1542–54.

17. Hukelmann JL, Anderson KE, Sinclair LV, Grzes KM, Murillo AB, Hawkins PT, et al. The cytotoxic T cell proteome and its shaping by the kinase mTOR. Nat Immunol. 2016;17(1):104–12.

18. Marchingo JM, Sinclair LV, Howden AJ, Cantrell DA. Quantitative analysis of how Myc controls T cell proteomes and metabolic pathways during T cell activation. Elife. 2020;9.

19. Tan H, Yang K, Li Y, Shaw TI, Wang Y, Blanco DB, et al. Integrative Proteomics and Phosphoproteomics Profiling Reveals Dynamic Signaling Networks and Bioenergetics Pathways Underlying T Cell Activation. Immunity. 2017;46(3):488–503.

20. Rieckmann JC, Geiger R, Hornburg D, Wolf T, Kveler K, Jarrossay D, et al. Social network architecture of human immune cells unveiled by quantitative proteomics. Nat Immunol. 2017;18(5):583–93.

21. Pircher H, Bürki K, Lang R, Hengartner H, Zinkernagel RM. Tolerance induction in double specific T-cell receptor transgenic mice varies with antigen. Nature. 1989;342(6249):559–61.

22. Nie Z, Hu G, Wei G, Cui K, Yamane A, Resch W, et al. c-Myc is a universal amplifier of expressed genes in lymphocytes and embryonic stem cells. Cell. 2012;151(1):68–79.

23. Hughes CS, Foehr S, Garfield DA, Furlong EE, Steinmetz LM, Krijgsveld J. Ultrasensitive proteome analysis using paramagnetic bead technology. Mol Syst Biol. 2014;10(10):757.

24. Sinclair LV, Howden AJ, Brenes A, Spinelli L, Hukelmann JL, Macintyre AN, et al. Antigen receptor control of methionine metabolism in T cells. Elife. 2019;8.

25. Wisniewski JR, Hein MY, Cox J, Mann M. A “proteomic ruler” for protein copy number and concentration estimation without spike-in standards. Mol Cell Proteomics. 2014;13(12):3497–506.

26. Tyanova S, Temu T, Sinitcyn P, Carlson A, Hein MY, Geiger T, et al. The Perseus computational platform for comprehensive analysis of (prote)omics data. Nat Methods. 2016;13(9):731–40.

27. Davies SP, Reddy H, Caivano M, Cohen P. Specificity and mechanism of action of some commonly used protein kinase inhibitors. Biochem J. 2000;351(Pt 1):95–105.

28. Miyagawa F, Zhang H, Terunuma A, Ozato K, Tagaya Y, Katz SI. Interferon regulatory factor 8 integrates T-cell receptor and cytokine-signaling pathways and drives effector differentiation of CD8 T cells. Proceedings of the National Academy of Sciences of the United States of America. 2012;109(30):12123–8.

29. Rollings CM, Sinclair LV, Brady HJM, Cantrell DA, Ross SH. Interleukin-2 shapes the cytotoxic T cell proteome and immune environment-sensing programs. Sci Signal. 2018;11(526).

30. Intlekofer AM, Takemoto N, Wherry EJ, Longworth SA, Northrup JT, Palanivel VR, et al. Effector and memory CD8+ T cell fate coupled by T-bet and eomesodermin. Nat Immunol. 2005;6(12):1236–44.

31. Kaech SM, Cui W. Transcriptional control of effector and memory CD8+ T cell differentiation. Nature reviews Immunology. 2012;12(11):749–61.

32. MacIver NJ, Michalek RD, Rathmell JC. Metabolic regulation of T lymphocytes. Annu Rev Immunol. 2013;31:259–83.

33. Buck MD, O’Sullivan D, Pearce EL. T cell metabolism drives immunity. J Exp Med. 2015;212(9):1345–60.

34. Larange A, Cheroutre H. Retinoic Acid and Retinoic Acid Receptors as Pleiotropic Modulators of the Immune System. Annu Rev Immunol. 2016;34:369–94.

35. Coleman ML, Marshall CJ, Olson MF. Ras promotes p21(Waf1/Cip1) protein stability via a cyclin D1-imposed block in proteasome-mediated degradation. Embo j. 2003;22(9):2036–46.

36. Zannini L, Delia D, Buscemi G. CHK2 kinase in the DNA damage response and beyond. J Mol Cell Biol. 2014;6(6):442–57.

37. Alfei F, Kanev K, Hofmann M, Wu M, Ghoneim HE, Roelli P, et al. TOX reinforces the phenotype and longevity of exhausted T cells in chronic viral infection. Nature. 2019;571(7764):265–9.

38. Khan O, Giles JR, McDonald S, Manne S, Ngiow SF, Patel KP, et al. TOX transcriptionally and epigenetically programs CD8(+) T cell exhaustion. Nature. 2019;571(7764):211–8.

39. Utzschneider DT, Charmoy M, Chennupati V, Pousse L, Ferreira DP, Calderon-Copete S, et al. T Cell Factor 1-Expressing Memory-like CD8(+) T Cells Sustain the Immune Response to Chronic Viral Infections. Immunity. 2016;45(2):415–27.

40. Betts BC, Bastian D, Iamsawat S, Nguyen H, Heinrichs JL, Wu Y, et al. Targeting JAK2 reduces GVHD and xenograft rejection through regulation of T cell differentiation. Proceedings of the National Academy of Sciences of the United States of America. 2018;115(7):1582–7.

41. Valvezan AJ, Manning BD. Molecular logic of mTORC1 signalling as a metabolic rheostat. Nature metabolism. 2019;1(3):321–33.

42. Wang R, Dillon CP, Shi LZ, Milasta S, Carter R, Finkelstein D, et al. The transcription factor Myc controls metabolic reprogramming upon T lymphocyte activation. Immunity. 2011;35(6):871–82.

